# Colony-level pollen collection reflects visitation of managed bumble bees (*Bombus impatiens*) in strawberry fields and surrounding landscapes without reducing pollen limitation

**DOI:** 10.1101/2025.10.04.677857

**Authors:** Leeah I Richardson, Olivia Miller, David Sossa, Aaron Iverson, Scott McArt, Katja Poveda, Heather Grab

## Abstract

Managed bees are frequently used to supplement pollination of a range of crop plants. Yet, their effectiveness varies and much of their foraging can be outside of the focal crop. Bee foraging behavior is highly sensitive to the availability of alternative floral resources which depend on the composition of the surrounding landscape. To understand how foraging and crop pollination dynamics of the managed common eastern bumble bee (*Bombus impatiens*) are influenced by landscape composition, we assessed their visitation and pollination in cultivated strawberry fields along a landscape gradient. We also used pollen extracted from bumble bee (*Bombus impatiens*) colony wax to understand whole-colony foraging over the blooming period. Bumble bee visits to strawberry crop flowers were very low and pollen limitation was not reduced at sites with greater visitation; however, colonies in fields with higher visitation rates had more pollen from the Rosaceae family (including strawberry). We found that landscape composition affected foraging effort on strawberry at the colony level, as a lower proportion of rosaceous pollen was collected in landscapes with more surrounding pasture, while a greater proportion of rosaceous pollen was collected at sites with larger strawberry fields. Overall, our results suggest that managed bumble bees do forage on strawberry crop flowers, but they predominantly forage elsewhere, especially in surrounding pastures when strawberry fields are small. Therefore, the addition of bumble bee colonies may not be economically beneficial for strawberry growers at least in diverse agricultural landscapes similar to that of our study region.

## Introduction

Pollinators serve an important role in food production, with about 30% of agricultural food production relying to some extent on animal mediated pollination (Klein *et al*., 2007; Khalifa *et al*., 2021). Yet, agriculture itself is a major driver of pollinator decline (Cane & Tepedino, 2001) due to factors like habitat loss and exposure to pesticides (Goulson *et al*., 2015) leading to reductions in the services provided by wild bees frequently observed in landscapes that are more agriculturally dominated (Ricketts *et al*., 2018; Grab *et al*., 2019). To achieve sufficient pollination, growers can bring in managed bees to supplement natural pollination (Rucker *et al*., 2012). However, supplemental managed bees may not always improve yields (Petersen *et al*., 2013) and factors that moderate their efficacy are unclear.

Supplementing crop pollination with managed pollinators does not always increase pollination rates in the focal crop (Petersen *et al*., 2013; Trillo *et al*., 2018) for two potential reasons. First, in landscapes with high amounts of natural habitat, wild pollinators are often more abundant and more likely to meet crop pollination demands on their own (Garibaldi *et al*., 2011; Kennedy *et al*., 2013; Mallinger & Gratton, 2015). Second, in environments with abundant natural habitats, managed pollinators may preferentially forage in these areas and overlook the focal crop (Guzman *et al*., 2019). For example, in cranberry agroecosystems honey bee stocking density was found to have a strong positive relationship with yield in fields that have low forest cover but no relationship to yield in landscapes with high forest cover (Gaines-Day & Gratton, 2016), where around only 15% of pollen from hives was from cranberry (Guzman *et al*., 2019). These low rates of focal crop pollen collection by managed pollinators have been reported from a range of other crop systems (8.7% apple (McArt *et al*., 2017a), 0.4-2% pumpkin (Brochu *et al*., 2020), 19-26% strawberry (Bänsch *et al*., 2020), 0.5-26% blueberry (Graham *et al*., 2023)), and suggest that managed bees are frequently using resources outside of the focal crop. For these reasons, we expect managed bees to be more effective in simple, agriculturally dominated landscapes than in landscapes containing more natural and semi-natural habitats.

The honey bee (*Apis mellifera*) is not the only managed bee that can be used to supplement pollination services (Osterman *et al*., 2021). The common eastern bumble bee (*Bombus impatiens*) is widely used across the eastern United States as a commercially available alternative to honey bees for supplemental pollination services, as they are generalist foragers that are capable of pollinating many different crops (Evans, 2017). While these colonies are primarily used for pollination within greenhouses (Evans, 2017), they are also sold to supplement wild pollinators for crops grown in fields (Stanghellini et al., 1998; Desjardins & De Oliveira, 2006; Artz & Nault, 2011). Though bumble bee colonies have fewer individuals than honey bees, they can be more effective at pollinating certain crops (ex. blueberry, tomatoes, etc) due to traits like hairiness (pilosity) (Stavert et al., 2016), tolerance to foraging in colder temperatures (Corbet *et al*., 1993), and the ability to buzz pollinate <u>(De Luca & Vallejo-Marín, 2013)</u>. They can also have a higher per-visit efficiency and colonies (which only live one season) can be purchased for prices comparable to or even less expensive than renting commercial honey bees for one season of pollination <u>(Drummond, 2012)</u>.

Strawberries are a crop that has been shown to benefit from bumble bee pollination in greenhouses and other enclosed environments <u>(Dimou *et al*., 2008</u>; <u>Martin *et al*., 2019)</u>. However, the majority of strawberry production occurs in open field conditions <u>(*USDA NASS, 2022*)</u>. Insect pollination generally benefits strawberry crops (Gudowska *et al*., 2024; Menzel, 2023). However, adding managed honey bee colonies to strawberry plantings has been found to negatively affect yield (Angelella et al. 2021), while increased wild bee abundance is associated with improved yield <u>(Connelly *et al*., 2015)</u>. In our study region, the most common pollinators to strawberry during the early spring are bees in the genus *Andrena, Lasioglossum*, and *Augochlorella;* followed by honey bees, which were found to make up less than 7% of the pollinator community (Connelly *et al*., 2015). This is similar to the composition detected in another study region, also in eastern North America, although it was less dominated by honey bees (MacInnis & Forrest, 2019). Bumble bees are considered effective pollinators generally based on their size and loosely packed pollen (Nye & Anderson, 1974), while managed bumble bees (*B. terrestris*) have been shown to dedicate much of their foraging effort to strawberry plantings in Spain (Trillo *et al*., 2020).

In the northeast United States, wild bumble bee foundress queens begin to start colonies during strawberry bloom from May to early June, but are not established to produce foraging workers until after the bloom of spring strawberry has completed (Lanterman *et al*., 2019). Indeed, previous work suggests that bumble bees do not visit spring strawberry plantings in New York at sites where they are not actively managed (Connelly *et al*., 2015). This suggests the potential for commercially available bumble bee colonies to provide supplemental services, particularly in agriculturally dominated landscapes where pollination services from wild bees may be lower (Connelly *et al*., 2015).

Understanding which floral resources bumble bees are foraging on by identifying pollen grains offers an important insight into their foraging patterns (Guzman *et al*., 2019). It is typically difficult to get this colony level information because floral usage in bumble bees varies between individuals (Saifuddin & Jha, 2014), across the season (Wood *et al*., 2018), and even over the course of a single day (Inouye, 1978). This means that short sampling periods are unlikely to capture the true breadth of floral resource use, and it is typically impractical to look at the pollen loads of every returning forager to a colony. However, techniques can be adapted for extracting pollen from beeswax (Furnessa, 1994) to look at the pollen brought back at the colony level, which allows integration over time and across individuals with a single sample from a colony. These methods have recently been adapted to look at the pollen found in medieval wax seals (Kasso *et al*., 2023), and to understand historic honey bee foraging patterns from museum comb specimens (Kasso *et al*., 2024); their use in investigating variation in contemporary bumble bee foraging offers a promising new direction to understanding season-long colony level foraging efforts.

In this study, we assessed the degree to which supplemental managed colonies of *B. impatiens* visit strawberry plantings in open field settings during peak bloom. Specifically, we focus our analyses on four questions: (1) Does managed bumble bee visitation rate to strawberry depend on surrounding land cover? (2) Do fields with greater visitation by managed bumble bees have reduced pollen limitation? (3) Is visitation rate by managed bumble bees reflected in colony Rosaceae pollen collection? and (4) Is pollen collection by managed bumble bees is dependent on landscape context?

## Methods

### Landscape composition and colony placements

In the spring of 2018, we surveyed 10 strawberry fields across upstate New York (Fig. 1A) that were a minimum of 500 m apart, placing commercially available bumble bee (Bombus impatiens) colonies from Koppert Biological Systems (MI, USA) in or along the edge of each field. Colonies were placed in the fields between May 14–15^th^ (all within a 24 hour period), just prior to strawberry bloom, and remained in the fields until fruit had set in mid June. Fields ranged in size from 0.069 hectares to 3.035 hectares. The number of colonies placed in each field ranged from 1–16 depending on field size, such that stocking densities were maintained at approximately two colonies per acre (0.4 ha) as recommended by Koppert Biological Systems, for a total of 43 colonies across all sites.

**Figure 1:**
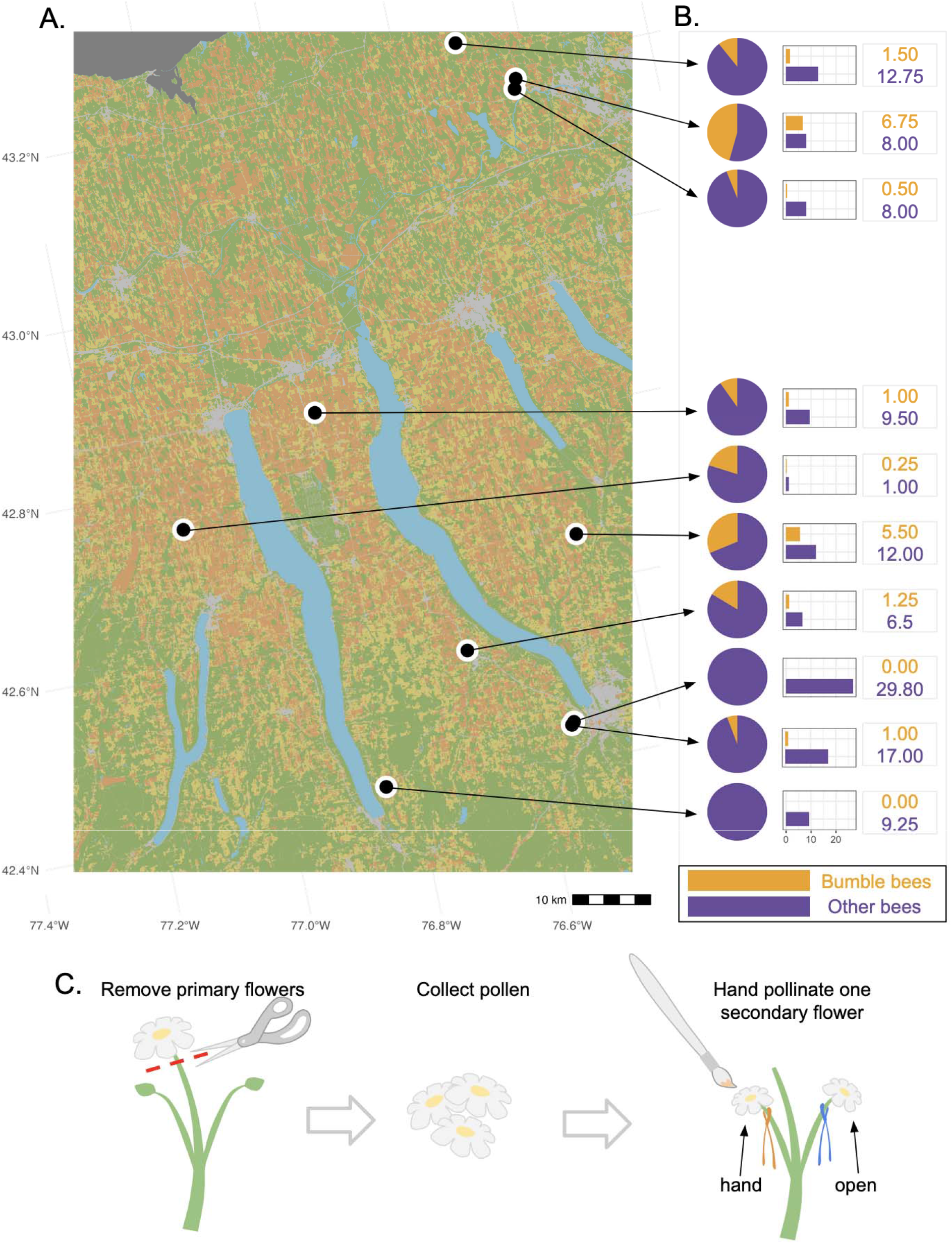
**A)** Site map depicting 2018 landscape categories by color (brown indicating agricultural land cover, yellow indicating pasture land cover, and green indicating semi-natural land cover). **B)** Charts showing the proportion and number of observed bee visits to strawberry flowers by bumble bees vs all other bees at each of the 10 fields. The average number of visits to strawberry flowers by bumble bees (shown in yellow) and by other bees (shown in purple) are also displayed for each site. **C)** Method of collecting pollen from removed primary flowers to hand-pollinate one of the secondary flowers while leaving the other to be open-pollinated in the fields to assess the influence of bee visitation on pollination limitation.

We determined the landscape composition around each field using the 2018 Cropland Data Layer (https://nassgeodata.gmu.edu/CropScape/). The proportion of each land cover category was calculated at 500, 750 and 1000 m through ArcGIS 10.3 (ESRI 2018) for three different land cover types: agricultural (cultivated crops; ranging from 5.9%-86.1% at 1000 m), semi-natural (forests, shrublands, wetlands; ranging from 2.4%-24.4% at 1000 m), and pasture/grass/hay (alfalfa, meadows, clover, wildflowers, grasses; ranging from 5.8% to 30.0% at 1000 m), hereafter referred to as ‘pasture’ (Figure 1).

### Bumble bee visitation

We conducted visual surveys to determine the visitation rate of bumble bees to strawberry during the blooming period. Four times during strawberry bloom (May 16th - June 7th) we walked a 10 min transect in each field at a consistent slow pace and counted each bee that visited a flower. Visits were defined as contact with the flower petals or stigmas (Grab *et al*., 2018), and we visually identified each bee as either a bumble bee or any other type of bee (clade: Anthophila), which were primarily bees in the genera *Andrena, Lasioglossum, Augochlorella*, and *Apis*. Because wild bumble bee workers were not yet foraging in this region during this time period (Lanterman *et al*., 2019) and previous studies in the region reported no bumble bee visitors to strawberry at fields without managed colonies (Connelly *et al*., 2015), we assumed that all bumble bee visits during our sampling periods were from our managed *B. impatiens* colonies.

### Sentinel plants for assessing pollination

We started bare-root strawberry plants of the cultivar ‘Seascape’ (Indiana Berry, IN, USA) in 10 cm pots in a greenhouse approximately 5 weeks before the onset of spring field strawberry bloom to use as sentinel plants flowering simultaneously with field-grown plants. Most of our 10 sites received 24 plants, although one site only received 18. Strawberry flowers tend to bifurcate from the stem internodes such that one large “primary” flower head is on the same stem as two smaller paired “secondary” flowers that typically open on the same day and will lead to fruits of similar sizes (Petran & Hoover, 2018). We removed the primary flowers from our plants, leaving the paired secondary flower buds, and dried the anthers from these primary flowers until they dehisced. On the day of secondary flower anthesis we used pollen collected from primary flowers to hand-pollinate one of the secondary flowers by gently brushing pollen across all stigmatic surfaces with a small paint brush (hand-pollinated) while we left the other secondary flower to be pollinated in the field (open-pollinated). These were marked with a colored piece of yarn tied loosely around the base of each flower (Fig. 1C).

Plants were brought out to the fields on the same day that they were hand-pollinated and placed in trays that held ∼3 cm of water in groups of 6 within the field rows when the field was flowering. Sentinel plants were only being open-pollinated when the field strawberries were also blooming and were present throughout the blooming period as we brought out trays on four dates throughout the season (first trays of plants to fields from May 14th - May 25th and the final plants were brought to fields May 31st - June 7th). We watered the plants during all field visits, and brought them back to be cared for in the greenhouse when the flowers had senesced (June 6th - June 22nd). Once the fruits had fully developed (June 7th - July 5th) we weighed each as a measure of pollination quality (Gudowska et al., 2024), and calculated the degree of pollen limitation in the field as the weight of the open-pollinated fruit subtracted from the hand-pollinated fruit then divided by the weight of the hand-pollinated fruit for each plant (Larson & Barrett, 1999; Baskin & Baskin, 2018). This allowed us to understand how close the open-pollinated fruits were to being fully pollinated (represented by the hand-pollinated treatment) to quantify pollen limitation in this system. If we detected no difference between these treatments it would indicate that pollination was sufficient, due to animal-mediated pollination and/or self pollination.

We were able to retrieve paired fruits from 154 plants for an average (+/-standard deviation) of 15.40 +/-4.64 plants per field. Plants or their fruits that were lost were due primarily to herbivory or disease, which was distributed fairly evenly across fields (75 total plants were excluded from analyses: average 7.5 plants +/-4.67 SD per site).

### Wax and pollen extraction

At the end of strawberry bloom, we removed approximately 10 g of wax from a single randomly selected representative colony from each field to quantify pollen composition such that each site was represented by one colony. Because each colony grows over the course of the season, old wax is located in the center of the colony (Sakagami *et al*., 1967) and is much darker in color (Michener, 1974) so it is possible to only remove newer wax that had built up along the entire periphery after the colony was put into the strawberry fields. We expected the pollen present in the wax to be representative of pollen collection over time by random contamination into the wax during larval feeding and wax manipulation for construction (Hines *et al*., 2007). We pulverized the wax samples from these representative colonies in liquid nitrogen with a mortar and pestle for homogenization and to maximize the available surface area for pollen extraction.

We used a small sample (∼0.5 g) of this pulverized wax to undergo an acetolysis protocol to extract and identify the pollen that was present. We separated the pollen grains from the wax by adding 1 mL of glacial acetic acid to the wax sample in a 2 mL centrifuge tube and allowing it to soak for 24 hours so that pollen grains entered solution (Furnessa, 1994). This solution was then drained from the wax residue and centrifuged for 6 min at 13,000 rpm to form a pollen pellet.

A standard acetolysis protocol was then carried out as follows (*sensu* Jones, 2014): we drained the remaining acetic acid and then added 1 mL of a 9:1 solution of acetic anhydride and sulfuric acid and vortexed each tube. These samples were immediately placed into a 5 min hot water bath at 90°C and were then centrifuged for 5 min at 13,000 rpm. We removed the acetic anhydride and sulfuric acid solution from the pellet and added a 50% ethanol solution to rinse away any remaining solution. Samples were vortexed again and then underwent a final 5 min centrifuge at 13,000 rpm. The pellet was again rinsed with the 50% ethanol solution and then gently mixed with a small volume of glycerol to store the pollen grains for later identification.

### Pollen counting and identification

After a minimum of 24 hours we added 15 uL of this glycerol solution and a small volume of fuschin stain to slides, and sealed the cover slip to the slide with clear nail polish. For consistency, we counted pollen at 400x magnification along horizontal transects, starting at the midline of the slide and establishing further transects bisecting the midline and slide edge until the first 300 pollen grains were counted to systematically quantify the composition of pollen collected by a colony (Jones & Bryant, 1998). A pollen grain was only counted if it was entirely visible in the field of view, and all pollen grains visible in the field of view once a count of 300 was achieved were counted (for a final count of 300-306 per slide). Pollen was identified as being within the family Rosaceae (Kapp *et al*., 2000), or was categorized as non-rosaceous.

### Statistical analyses

All analyses were done with R version 4.4.2 (R Core Team, 2024), and checked for violations of model assumptions (package: *DHARMa*; Hartig 2024). To test if bumble bee visitation rates differed from visitation rates by all other bees to strawberry flowers we used a linear mixed effect model (package: *lme4*; Bates *et al*., 2015) to assess if the average number of bees recorded was predicted by type of bee (bumble bee vs other bees) with field site as a random factor.

#### 1. Does managed bumble bee visitation rate to strawberry depend on surrounding land cover?

We evaluated if landscape composition surrounding each field influenced bumble bee visitation to strawberry flowers by running three individual linear models, one for each land cover category (agriculture, pasture, and semi-natural) as a predictor with the transformed (log + 1) average number of bumble bee visits as our response variable. We also evaluated if field size influenced bumble bee visitation with linear models. The average number of bumble bee visits was log + 1 transformed in all analyses to aid in model fit. We estimated predictor significance with ANOVA likelihood ratio tests. We use the landscape 1000 m surrounding the fields because this scale has previously been shown to be most relevant to bees foraging on strawberry in this region (Connelly et al. 2015) and relevant to bumble bee foraging distances (mean: 842 m (McArt *et al*., 2017b)), but also evaluated the 750 m and 500 m scales (Table 1).

**Table 1:** Results from models across three spatial scales (1000 m, 750 m, and 500 m) for landcover categories. Significant effects are bolded. See visualizations of effects at the 1000 m scale in supplementary information (Fig. S1).

| Response | Landcover Category | Surrounding Scale | Test Statistic | P-Value |
| --- | --- | --- | --- | --- |
| Bumble bee visitation<br>(log + 1 transformed) | Pasture | 1000 m | F = 1.043 | p = 0.337 |
|  |  | 750 m | F = 0.476 | p = 0.537 |
|  |  | 500 m | F = 0.414 | p = 0.537 |
|  | Agriculture | 1000 m | F = 0.002 | p = 0.968 |
|  |  | 750 m | F = 0.001 | p = 0.972 |
|  |  | 500 m | F = 0.011 | p = 0.918 |
|  | Semi-Natural | 1000 m | F = 0.689 | p = 0.431 |
|  |  | 750 m | F = 0.692 | p = 0.429 |
|  |  | 500 m | F = 0.077 | p = 0.788 |
| Proportion<br>rosaceous pollen | Pasture | **1000 m** | **z = -2.092** | **p = 0.036** |
|  |  | 750 m | z = -1.634 | p = 0.102 |
|  |  | 500 m | z = -1.599 | p = 0.109 |
|  | Agriculture | 1000 m | z = 0.846 | p = 0.397 |
|  |  | 750 m | z = 0.827 | p = 0.408 |
|  |  | 500 m | z = 1.060 | p = 0.289 |
|  | Semi-Natural | 1000 m | z = -1.540 | p = 0.123 |
|  |  | 750 m | z = -1.433 | p = 0.151 |
|  |  | 500 m | z = -0.651 | p = 0.514 |
| Pollen limitation:<br>(hand - open) / hand | Pasture | 1000 m | $X^2 = 1.266$ | $p = 0.260$ |
| | | 500 m | $X^2 = 0.776$ | $p = 0.378$ |
| | | 750 m | $X^2 = 0.566$ | $p = 0.451$ |
| | Agriculture | 1000 m | $X^2 = 2.996$ | $p = 0.083$ |
| | | 750 m | $X^2 = 2.412$ | $p = 0.120$ |
| | | 500 m | $X^2 = 1.745$ | $p = 0.186$ |
| | Semi-Natural | 1000 m | $X^2 = 2.372$ | $p = 0.123$ |
| | | 750 m | $X^2 = 2.427$ | $p = 0.119$ |
| | | 500 m | $X^2 = 1.754$ | $p = 0.185$ |

#### 2. Do fields with greater visitation by managed bumble bees have reduced pollen limitation?

To then assess if fields with greater observed bee visitation had reduced pollen limitation we ran a linear mixed effects model (package: *lme4*; Bates *et al*., 2015) with the difference between the open and hand-pollinated fruit divided by the hand-pollinated fruit as the response and the transformed (log + 1) average number of bumble bee visits as the predictor, we also ran these models with average number of other bee visits (log + 1) and with average total number of bee visits (log + 1) as predictors to understand the role of increased visitation by all bees. Last, we ran a model with field size as a predictor to understand the role that the amount of cultivated strawberry around the colonies had on pollen limitation. We included site as a random factor to account for the multiple plants from each field. We estimated predictor significance with ANOVA likelihood ratio tests.

#### 3. Is visitation rate by managed bumble bees reflected in colony Rosaceae pollen collection?

To understand the relationship between observed bumble bee visitation at our focal crop and the proportion of rosaceous pollen we recovered from the colonies we ran a beta regression model (package: *betareg*; Cribari-Neto & Zeileis, 2012) with the transformed (log + 1) average number of bumble bee visits as the predictor and the proportion of rosaceous pollen as the response. We estimated predictor significance for the beta regression with a likelihood ratio test (package: *lmtest;* Zeileis & Hothorn, 2002).

#### 4. Is pollen collection by managed bumble bees dependent on landscape context?

We then evaluated if landscape composition surrounding each field influenced the proportion of rosaceous pollen from the colony by running three models (package: *betareg*; Cribari-Neto & Zeileis, 2012), one for each land cover category (agriculture, pasture, and semi-natural) as a predictor with the proportion rosaceous pollen as the response variable. We also ran a model with field size as a predictor to understand the role that the amount of cultivated strawberry around the colonies had on rosaecous pollen collection. As above, we estimated predictor significance with likelihood ratio tests (package: *lmtest;* Zeileis & Hothorn, 2002).

## Results

Across the 4 visitation surveys to the 10 sites we observed a total of 648 bees visiting strawberry flowers: 72 bumble bees, and 576 other bees. Bumble bees provided substantially fewer visits to strawberry flowers compared to other bees (Figure 1B; Χ^2^ = 12.786, df = 1, p < 0.001). Average bumble bee visits in the 10 min observation period across the 4 rounds per site ranged from 0 to 6.75 bees (mean = 1.78) while visits from other bees ranged from 1 to 26.80 bees (mean = 11.08).

### 1. Does managed bumble bee visitation rate to strawberry depend on surrounding land cover?

The average number of bumble bee visits was not affected by surrounding pasture (F_1,8_ = 1.043, p = 0.337), semi-natural area (F_1,8_ = 0.689, p = 0.431), or agriculture (F_1,8_ = 0.002, p = 0.968) at any scale (Table 1). Strawberry field size also did not affect bumble bee visitation (F_1,8_ = 0.983, p = 0.350).

### 2. Do fields with greater visitation by managed bumble bees have improved pollination?

There was no evidence that managed bumble bees improved pollination to strawberry as sites with greater bumble bee visitation did not have reduced pollen limitation (Figure 2B; X^2^ = 0.099, df = 1, p = 0.754); increased visitation by other bees also did not have an effect (X^2^ < 0.001, df = 1, p = 0.979), nor did increased total visitation (X^2^ = 0.194, df = 1, p = 0.659). We also did not find that surrounding landscape (1000 m surrounding the site) or field size affected pollen limitation (pasture: X^2^ = 1.266, df = 1, p = 0.260; agriculture: X^2^ = 2.996, df = 1, p = 0.083; semi-natural area: X^2^ = 2.372, df = 1, p = 0.123; field size: X^2^ = 0.877, df = 1, p = 0.348) at any spatial scale (Table 1).

**Figure 2:**
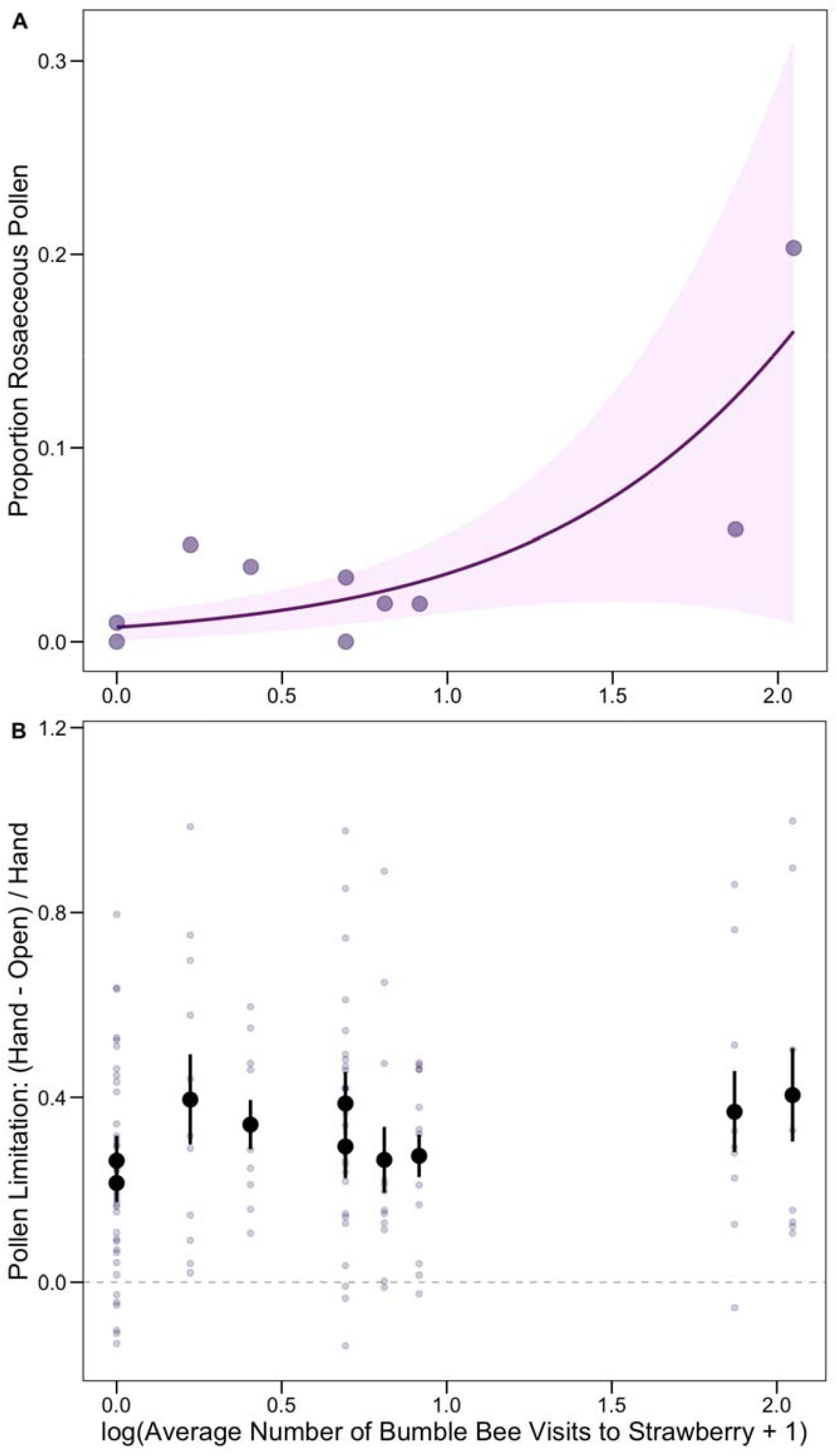
Increased bumble bee visitation led to more rosaceous pollen in the colony wax but did not affect pollen limitation. **A:** Relationship between the transformed average number of bumble bee visits to strawberry flowers observed across the 4x 10 min visitation surveys and the proportion of rosaceous pollen recovered from the colony wax with beta-regression 95% confidence intervals. **B:** Pollen limitation data with each point representing the value calculated from the masses of a pair of fruit from a single plant ((Fruit_hand_ - Fruit_open_) / Fruit_hand_). The mean +/-SE for each site is depicted in black. The grey dotted line at zero represents if there was no difference between the hand-pollinated and open-pollinated fruit, all values above zero show pairs where the hand-pollinated fruit was larger than the open-pollinated fruit.

### 3. Is visitation rate by managed bumble bees reflected in colony Rosaceae pollen collection?

In line with the low visitation rates by bumble bees to strawberry, we recovered correspondingly low proportions of rosaceous pollen from the colony wax, ranging from 0 to 0.20 of total pollen grains screened (mean proportion ± sd= 0.04 ± 0.059). However, fields with more observed bumble bee visits to strawberry flowers had a greater proportion of rosaceous pollen (Figure 2A; z = 3.272, p = 0.001).

### 4. Is pollen collection by managed bumble bees dependent on landscape context?

Colonies at sites with larger strawberry fields collected more rosaceous pollen (Figure 3A; z = 3.878, p < 0.001). Bumble bee colonies collected less rosaceous pollen when there was more pasture in the surrounding landscape at a 1000 m radius (Figure 3B; z = -2.092, p = 0.036). The proportion of rosaceous pollen was not influenced by surrounding semi-natural area (z= -1.540, p = 0.123) or by surrounding agriculture (z = 0.846, p = 0.397) at any scale (Table 1).

**Figure 3:**
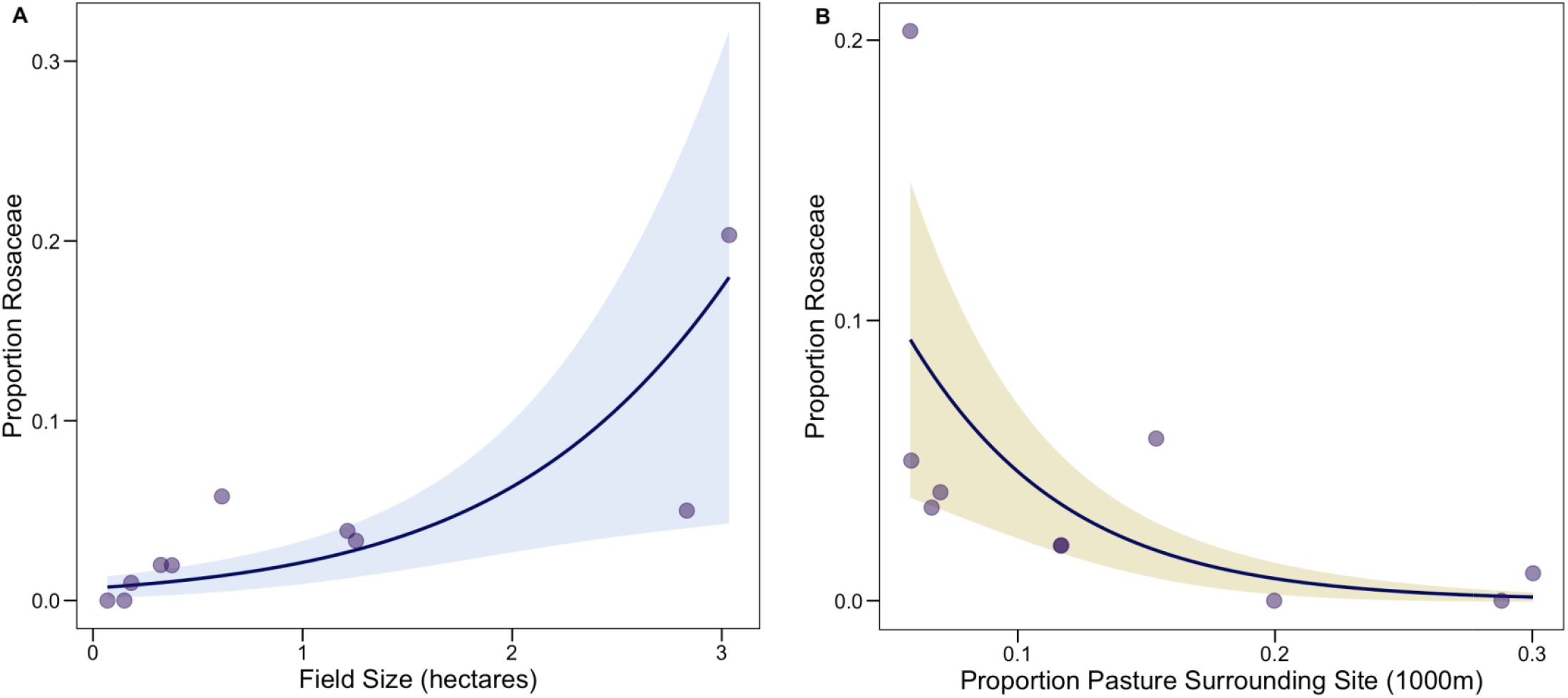
More rosaceous pollen was recovered from colony wax when strawberry fields were larger, and less rosaceous pollen was recovered when there was more pasture in the surrounding landscape. **A:** Relationship between strawberry field size (hectares) and the proportion of rosaceous pollen recovered from the colony wax with beta-regression 95% confidence interval. **B:** Relationship between the proportion of pasture surrounding 1000m around fields and the proportion of rosaceous pollen recovered from the colony wax with beta-regression 95% confidence interval.

## Discussion

Landscape composition is known to influence crop visitation by wild bees (Kennedy *et al*., 2013) but the role of landscape in mediating the efficacy of managed bees is less well understood. Here, we address this gap using a novel approach that infers whole-colony foraging by extracting pollen from nest wax. We find that managed bumble bees (*B. impatiens*) provided a relatively low number of visits to our focal strawberry crop (family: Rosaceae) compared to other bees, collected a low proportion of rosaceous pollen, and increased visitation by bumble bees did not reduce pollen limitation. These findings improve out knowledge of how managed bumble bees can be effectively used in agroecosystems and suggest that supplementing strawberry pollination with managed bumble bee colonies is not likely to improve pollination, particularly in landscapes where other resources found in pasture habitats may be more attractive to bumble bees, although future work could directly compare pollen limitation between sites with and without managed bumble bees.

### 1. Does managed bumble bee visitation rate to strawberry depend on surrounding land cover?

While observed bumble bee visitation was not influenced by landscape context, their collection of rosaceous pollen at the colony level was both positively correlated with focal crop visitation and field size, and lower at fields with more pasture in the surrounding landscape. This indicates that although there may not be fewer bumblebees (*B. impatiens)* visiting strawberry flowers as pasture increases, the colony overall is dedicating less of its foraging efforts toward the focal strawberry crop when these other resources are present. Similarly, while we did not observe more bumble bees on strawberry flowers when fields were larger, the colony seems to have been dedicating relatively more of its foraging effort toward strawberry, which could be because a greater proportion of their foraging radius is occupied by strawberry as they are central place foragers (Bell 1990). Future work could use a similar design but identify all pollen grains to species instead of just categorizing as either rosaceous or not to determine which floral resources compete the most with strawberry for bumble bee visits since bumble bees are known to have strong floral preferences (Lanterman Novotny *et al*., 2023).

### 2. Do fields with greater visitation by managed bumble bees have improved pollination?

Managed bumble bees in this system appeared to be primarily visiting other plants across our 10 sites since an average of only 4.3% of the pollen that these colonies collected was from plants within the family Rosaceae. This is counter to their intended purpose as supplemental pollinators to strawberry placed in the fields, but corresponds with work in other systems. For instance, managed honey bees collect less focal cranberry pollen in landscapes with high woodland cover which presumably contains preferred floral resources (Guzman *et al*., 2019). Bumble bees are known to have strong foraging preferences based both on floral traits (Harder, 1988), resource availability (Aleixo *et al*., 2017), and nutritional needs of the colony (Vaudo *et al*., 2016). When managed bumble bee colonies are added to squash fields less than five percent of the pollen collected is from squash (Brochu *et al*., 2020), and managed colonies are not effective at providing supplemental pollination services for squash, showing no increase in the amount of fruit produced per plant (Petersen *et al*., 2013). Similarly, we also find that greater visitation by bumble bees to strawberry did not reduce pollen limitation. Our hand-pollinated fruits tended to be larger than those that were open-pollinated, which suggests that our sites were generally pollen limited and so there may not have been enough bee visitation at any site to see improved pollination which corresponds with our finding that total bee visitation was not related to improved fruit set. As our sites appeared to be generally pollen-limited (our open-pollinated fruits tended to be smaller than the hand-pollinated fruits), this suggests that animal-mediated pollination and/or self fertilization did not fully maximize pollination. Increased stocking density of managed bumble bees may improve pollination, as stocking rates twice our stocking density have been observed to improve pollination in blueberry (Drummond et al. 2012). This is suggested by our finding that colonies in larger fields collected more rosaceous pollen despite similar colony densities per acre, inconsistent with a scenario in which strawberry flowers were saturated by managed bees and thus limited in pollen availability.

### 3. Is visitation rate by managed bumble bees reflected in colony Rosaceae pollen collection?

In this study we adapted a method of extracting pollen from museum wax sculptures by soaking wax acetic acid prior to acetolysis (Furnessa, 1994) and studying historic honey bee foraging (Kasso *et al*., 2023, 2024) for use in a contemporary ecological context to understand bumble bee foraging efforts at the colony scale. It has recently been suggested that when senesced bumble bee colonies are incidentally found in the wild data should be collected on colony demographics, size, architecture and pollen stored in pots to gain a better understanding of these understudied aspects of bumble bee colonies across species (Smith *et al*., 2025), and we suggest that extracting pollen from the wax broadly could also lend useful insights to the floral resources used. Due to the habit of building outward from a central point as the bumble bee colony grows (Sakagami *et al*., 1967) it may also be possible to look at colony-level pollen collection over time. This could allow for broader studies of colony foraging dynamics and resource usage; including from species that are not commercially managed.

### 4. Is pollen collection by managed bumble bees dependent on landscape context?

Bee diet composition can be influenced by the surrounding landscape (Inês da Silva *et al*., 2024). For instance, landscapes with more semi-natural habitats and lower agricultural cover have been associated with greater diet diversity and lower collection of Rosaceae pollen in the solitary managed bee, *Osmia cornifrons* (Centrella *et al*., 2020). While we do not see an effect of semi-natural or agricultural landcover, we do find our colonies collecting less rosaceous pollen in landscapes with more surrounding pasture, which similarly suggests that crop pollen collection is influenced by what other resources are available in the surrounding landscape. For example, clover (*Trifolium* spp.) is a preferred floral resource for *Bombus impatiens* (Lanterman Novotny *et al*., 2023) and is included in our pasture areas, while our semi-natural areas were largely forests with primarily wind pollinated species blooming during our study period (Regal, 1982). Other rosaceous crops (apple and raspberry) and wild rosaceous plants (ex: *Crataegus* spp., *Rubus* spp., and *Pyrus* spp.) were present in low abundance around some fields, and as pollen was not identified to species it is possible that visitation to these plants contributed to the patterns that we find. For example, co-blooming apple is known to reduce visitation to strawberry during peak bloom and then increase visitation later in the season (Grab *et al*., 2017) so if other rosaceous plants in the landscape were co-blooming with strawberry there could be increased pollen collection from these plants while simultaneously influencing strawberry visitation. However, the link between strawberry visitation and field size to Rosaceae pollen suggests the rosaceous pollen is likely strawberry.

## Conclusions

There are potential costs associated with bringing in managed bees to supplement pollination. There is the financial cost of the bees themselves (Baylis *et al*., 2021), but also they can have negative effects on wild pollinators (Paini, 2004) that may already be providing these services (Garibaldi *et al*., 2014). Managed bees can outcompete wild bees (Mallinger *et al*., 2017), spread disease (Graystock *et al*., 2013, 2016; Martin *et al*., 2021), and can even function as ecological traps that directly lead to wild bumble bee queen mortality (Miller *et al*., 2023). Only assessing these risks to the pollinator community at a focal crop will poorly capture the true impact if they are largely foraging elsewhere.

To accurately assess the potential benefits of managed bees we need to understand whole colony foraging dynamics. Managed bumble bees are effective pollinators for many pollinator-dependent greenhouse crops as they are not accessible to wild pollinators (Velthuis & Doorn, 2006) and some field crops (Stubbs & Drummond, 2001; Drummond, 2012). We find that bumble bee colony-level foraging effort dedicated to our focal crop was generally very low and landscape-context dependent. Furthermore, we found that pollen limitation did not vary based on the observed bumble bee visitation rates; however, future work could expand to include sites without bumble bees present across a larger number of sites to better understand the role that bumble bees may play in strawberry pollination. By applying a novel method to extract pollen from colony wax, we quantify whole-colony bumble bee foraging and assess whether colonies preferentially forage on a focal crop and which wild pollinator communities they may impact.

## Acknowledgements

We thank the growers that allowed us access to their fields, as well as to Kass Urban-Mead and Diana Obregon for pollen staining advice, and to Nate Flicker for help with data collection. This work was supported by the USDA NIFA Hatch Grant NYC-139303, USDA NIFA 2018-08603 and Hatch Grant NYC-139440.

## Notes

**<u>Conflict of Interest statement:</u>** We have no conflict of interest to declare.

### Competing Interest Statement

The authors have declared no competing interest.

### Summary of Updates

results and discussion have been reorganized under the research questions as subheadings. small text edits to improve clarity have been made throughout

