## Supplementary material for "Colony-level pollen collection reflects visitation of managed bumble bees (*Bombus impatiens*) in strawberry fields and surrounding landscapes without reducing pollen limitation": Fig. S1

**Supplemental Material**

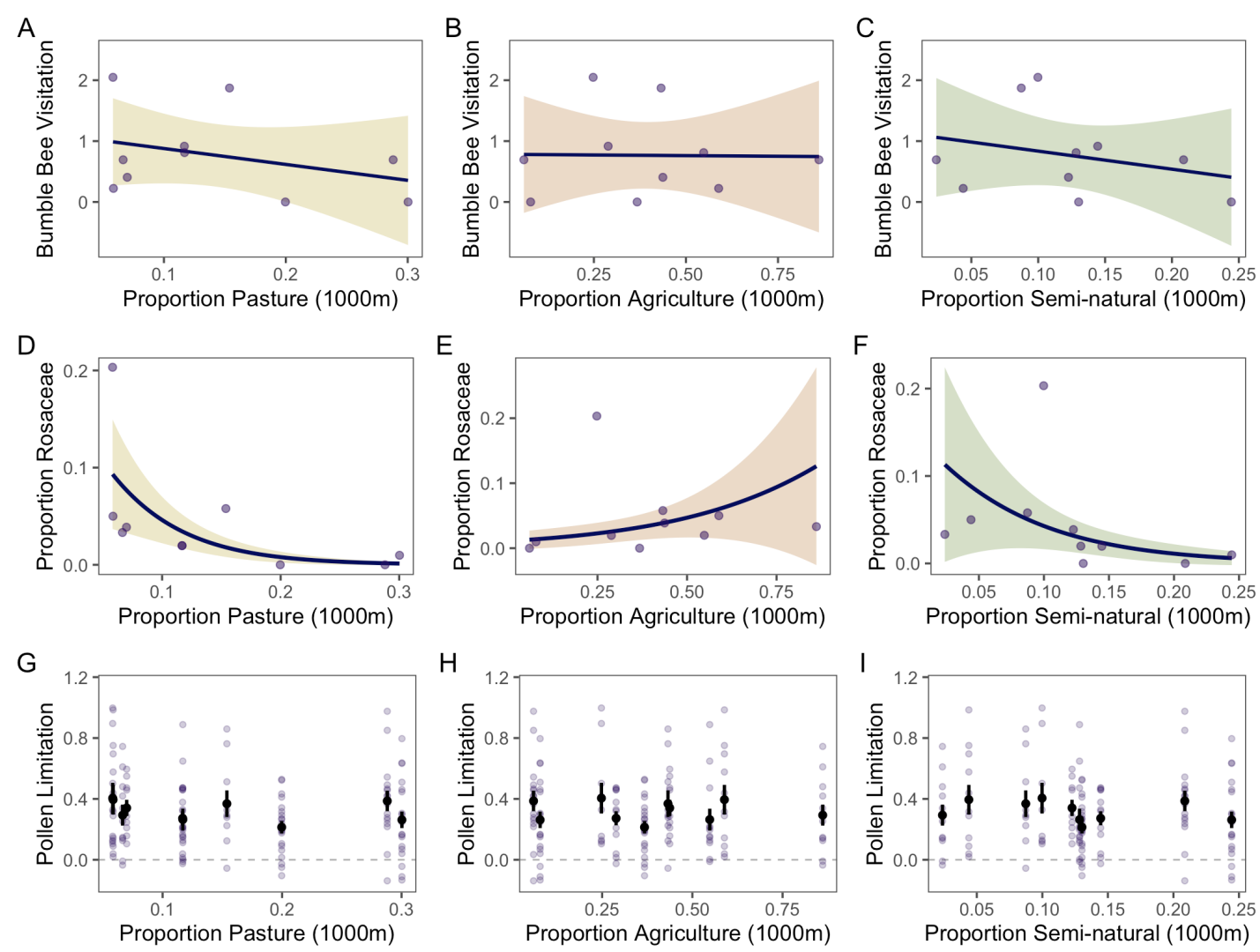

**Figure S1:** Relationships between landscape composition at 1000 m surrounding a site and response variables. **A.** Relationship between surrounding pasture and log(+1) transformed bumble bee visitation to strawberry. **B.** Relationship between surrounding agriculture and log(+1) transformed bumble bee visitation to strawberry. **C.** Relationship between surrounding semi-natural area and log(+1) transformed bumble bee visitation to strawberry. **D.** Relationship between surrounding pasture and proportion rosaceous pollen recovered from colony wax. **E.** Relationship between surrounding agriculture and proportion rosaceous pollen recovered from colony wax. **F.** Relationship between surrounding semi-natural area and proportion rosaceous pollen recovered from colony wax. **G.** Relationship between surrounding pasture and pollen limitation (hand - open)/hand. **H.** Relationship between surrounding agriculture and pollen limitation (hand - open)/hand. **I.** Relationship between surrounding semi-natural area and pollen limitation (hand - open)/hand.
